# Dissociation of disease phenotype and allele silencing in hypertrophic cardiomyopathy

**DOI:** 10.1101/642421

**Authors:** Alexandra Dainis, Kathia Zaleta-Rivera, Alexandre Ribeiro, Andrew Chia Hao Chang, Ching Shang, Feng Lan, Paul W. Burridge, Joseph C. Wu, Alex Chia Yu Chang, Beth L. Pruitt, Matthew Wheeler, Euan Ashley

## Abstract

Allele-specific RNA silencing has been shown to be an effective therapeutic treatment in a number of diseases, including neurodegenerative disorders. Studies of allele-specific silencing in hypertrophic cardiomyopathy to date have focused on mouse models of disease. Here, we investigate two methods of allele-specific silencing, short hairpin RNA (shRNA) and antisense oligonucleotide (ASO) silencing, using a human induced pluripotent stem cell-derived cardiomyocyte (hiPSC-CM) model of disease. We used cellular micropatterning devices with traction force microscopy and automated video analysis to examine each strategy’s effects on contractile defects underlying disease. We find that shRNA silencing ameliorates contractile phenotypes of disease, reducing disease-associated increases in cardiomyocyte velocity, force, and power. We find that ASO silencing, while better able to target and knockdown a specific disease-associated allele, showed more modest improvements in contractile phenotypes. We find a dissociation between allelic-specificity and functional improvements between the two tested therapeutic strategies, suggesting a more complex method of allelic control underlying HCM-associated transcripts.

**Author summary:** Allele-specific silencing, whereby a therapeutic molecule is used to lower the expression of just one of the two copies or alleles of a gene, may be a potential therapeutic strategy in diseases caused by a single mutation. In this paper, we examine two such strategies in hypertrophic cardiomyopathy, a disease characterized by an overgrowth of the left-ventricular heart muscle as well as contractile dysfunction. We used a human cell model of disease, creating induced pluripotent stem cell derived cardiomyocytes from a patient with HCM caused by a single base pair change in just one allele of the gene *MYH7*. We used two strategies to silence the disease-associated copy of *MYH7*, both focused on reducing RNA expression from the mutated allele, as well as state-of-the-art biophysical techniques for measuring contractility. We found that one silencing strategy, which reduced expression of both the disease-associated and the healthy alleles of *MYH7*, showed great improvements in contractility between treated and untreated cells. Our second strategy, which silenced only the disease-associated copy of *MYH7*, showed more modest improvements in contractility. This suggests that the disease mechanism underlying this type of hypertrophic cardiomyopathy may be more complex than just presence or absence of the mutated RNA.

## Introduction

RNA silencing has made recent strides towards clinical application, including antisense oligonucleotide therapeutics used in neurodegenerative disorders [1] and the recent approval of patisiran by the FDA as the first approved RNAi based drug [2]. These types of therapeutic strategies, which target disease at the RNA level, have the potential to be effective mechanisms of silencing disease-associated molecules before they can confer negative phenotypic effects, rather than treating symptoms after disease progression. One potential area where this type of strategy may be effective is in cardiovascular disease, including cardiomyopathies.

Hypertrophic cardiomyopathy, characterized by myocardial hypertrophy and disarray affects 1:500 in the population and can lead to atrial fibrillation, heart failure, and sudden death [3]. Causative genetic mutations can be identified in 30-60% of sequenced patients, and 75% of those appear in one of two genes: *MYH7* (myosin heavy chain 7) or *MYBPC3* (cardiac myosin binding protein c), key components of the cardiac sarcomere [4,5]. The first identified HCM-associated genetic mutation is a single base pair change that causes an arginine to glutamine change in the 403rd amino acid (*MYH7*-R403Q) [6]. Molecular analyses of this mutation have over time have revealed that it likely causes disease via a dominant-negative “poison-peptide” mechanism, whereby mutated protein products interfere with proper sarcomere function. Single-molecule, dual-beam optical trap studies of human myosin have found that the R403Q mutation causes a decrease in the intrinsic force of the myosin protein [7] while other studies, including examination of the effects of a small-molecule inhibitor of sarcomere power in mice, have indicated that the mutation results in an increase in sarcomere power output and hyperdynamic contraction [8]. Recent studies have provided additional insight into the potential mechanisms of *MYH7* based hypertrophic cardiomyopathy, including how mutations in *MYH7* may cause hypercontractility of the sarcomere by increasing the number of myosin heads available to interact with actin [9–11].

Allele-specific RNA silencing has shown therapeutic promise in scenarios where total gene knockdown may be detrimental. In these situations, targeted silencing of a disease-associated allele has shown potential to relieve disease phenotypes, including in some neurodegenerative diseases [12,13]. Allele-specific silencing has also shown promising results in mouse models of cardiovascular disease, including catecholaminergic polymorphic ventricular tachycardia [14] and HCM [15,16]. In humans, the dominant myosin isoform in the adult ventricle is β-myosin heavy chain, encoded by *MYH7*. Yet in the adult mouse, the dominant isoform is α-myosin heavy chain, encoded by *MYH6*. Murine studies of HCM have therefore focused on silencing and modulation of *MYH6* expression [15]. While informative, results of these studies can be hard to translate to potential human therapeutics, as α- and β-myosin have different contractile properties, and silencing of one does not necessarily recapitulate the phenotypic effects of silencing the other. For example, the same R403Q mutation in murine *MYH6* and *MYH7* showed different functional contractile consequences in in-vitro protein motility and enzymatic assays [17]. Though the R403Q mutation is one of the most studied mutations in HCM, the vast array of literature focused on the protein-level effects of this mutation have yet to agree on its effects on myosin function, likely due to the issues surrounding species, tissue, and isoform-level differences in myosin function, as well summarized by Nag et al. [7]. The genetic background in which these mutations are studied is therefore critical to appropriate interpretation of potential therapeutic strategies, and translational work must therefore aim to study the mutation in a model as close to the appropriate human genetic background as possible. One potential model for such studies is the use of patient-derived human induced pluripotent stem cells (hiPSCs).

Prior work in long QT syndrome has shown that allele-specific silencing of disease-associated mutations can rescue disease phenotypes in human induced pluripotent stem cell-derived cardiomyocytes (hiPSC-CMs) [18]. hiPSCs have become a widely used system for modeling a number of diseases in-vitro [19]. Their derivation to cardiomyocytes to study cardiovascular diseases, including hypertrophic cardiomyopathy, has allowed the field to study molecular and cellular phenotypes of disease, and test potential therapeutic interventions. This allows for investigation in a human cell context with an unlimited source of cells, rather than working in humanized animal models or with limited human biopsy tissue [20–25].

In order to appropriately study patient-derived hiPSC-CMs, assays have been developed to study multiple hypertrophy-associated phenotypes, including contractile defects. Traction force microscopy, which measures the forces cells apply to a substrate by measuring displacement of substrate embedded fiducials, has become a robust tool for quantifying force, velocity, and power in hiPSC-CMs, leading to discoveries about hiPSC-CM development, maturity, and response to environmental triggers [26–28]. More recent techniques have involved studying the contraction of cells using brightfield videos, removing the need for more extensive microscopic setups and allowing for the analysis of both individual cells and larger tissue-like structures [29]. These methods of studying contractility in our cell model system allow us to interrogate the effects of silencing at a single-cell level and gain more accurate insight into how novel therapeutics may ameliorate key disease phenotypes.

Here, we derive hiPSCs from siblings with severe HCM caused by the *MYH7*-R403Q familial mutation to create an in-vitro model of disease. We examine two potential methods of allele-specific gene silencing for both their ability to specifically silence a disease-causing allele in this model and for their ability to relieve disease phenotypes.

## Results

### hiPSC derivation from a family with multiple affected HCM patients

We derived hiPSCs and differentiated hiPSC-CMs from two members of a family with HCM. A family of five siblings with HCM showed signs of severe disease. Two of the five siblings passed away at young ages (34 and 45). The three remaining siblings all tested positive for the *MYH7*-R403Q mutation. One sibling suffered end stage heart failure and received a heart transplant at the age of 41. A more extensive family history revealed that the mother of the five siblings also had HCM, and died at the age of 73. While she did not undergo genetic testing, a number of additional early deaths in the maternal lineage (at the ages of 33, 39, 51, and 59) suggest penetrance across multiple generations. Our primary cell line was derived from a 45 year old female proband with the disease-associated *MYH7*-R403Q mutation (Fig 1A, patient 1, 52 years old at time of publication). The patient was diagnosed with HCM after presenting with a heart murmur at the age of 13. A second hiPSC cell line was derived by Cellular Dynamics Inc from an affected sibling (Fig 1A, patient 2).

**Fig 1:**
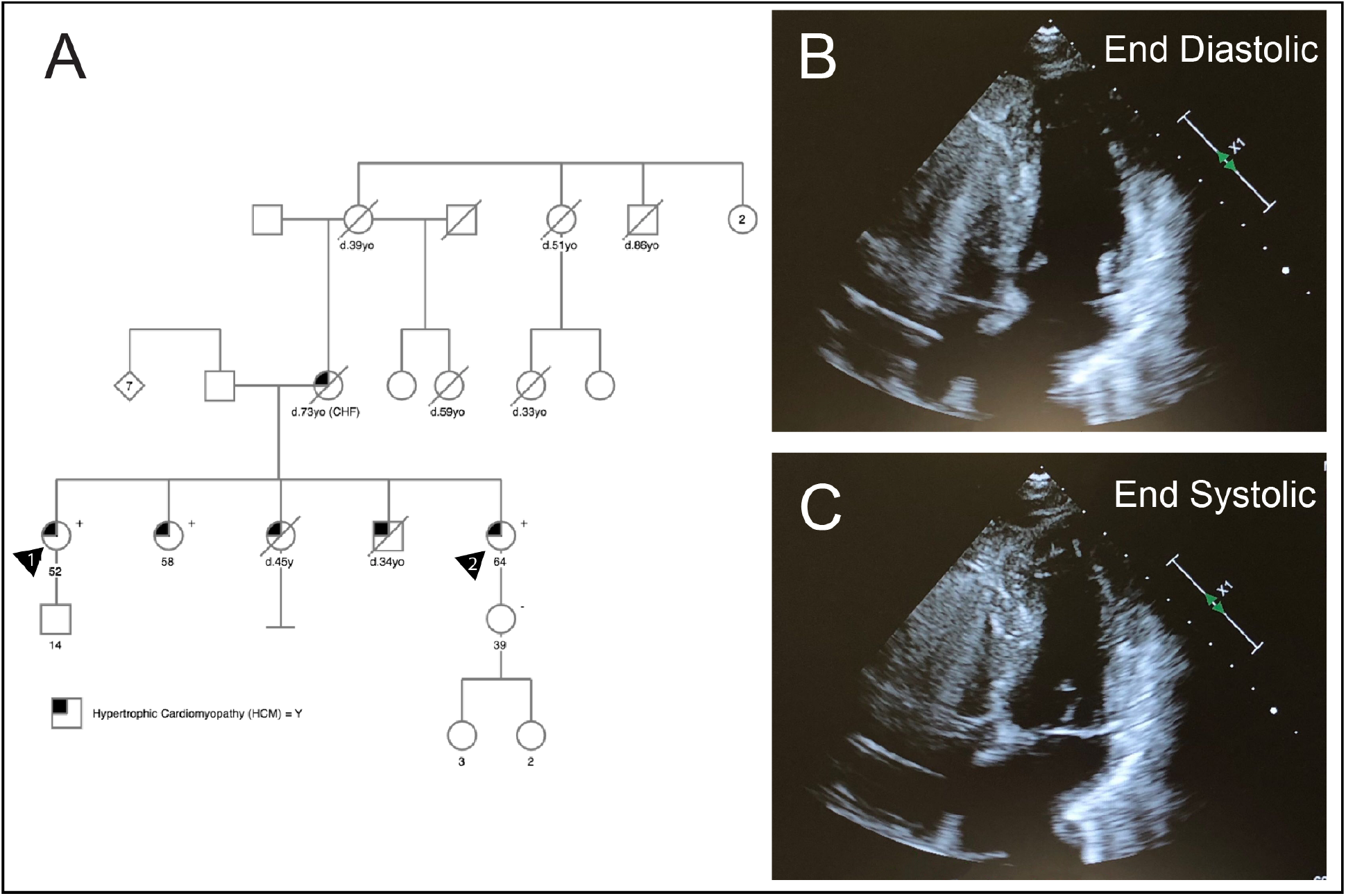
Family pedigree and hypertrophic phenotype of *MYH7*-R403Q HCM Patient. A) Family pedigree for patient 1 (designated by arrow 1) from whom the *MYH7*-R403Q hiPSC line was created. The patient and all four of her siblings showed symptoms of HCM. Two living siblings also tested positive for the R403Q mutation. One of the siblings (patient 2, designated by arrow 2) additionally donated cells for hiPSC derivation, which were used in the experiments in Fig 2C-G. B) and C) Apical four chamber echocardiography images of patient 1. Images demonstrate small cavity, left ventricular hypertrophy, and abnormal co-aptation of the mitral valve, as well as systolic anterior motion of the mitral valve.

### In-vitro shRNA silencing of R403Q allele in human hiPSC-CMs achieves knockdown and decrease in hypertrophic biomarkers

We used virally-delivered short hairpin RNA (shRNAs) to silence a dominant, HCM-associated mutation, *MYH7*-R403Q, in patient-derived hiPSC-CMs to show mutant allele knockdown in human cells. Potential shRNA designs were tested in a fluorescent-reporter HEK cell model to arrive at the mutant-allele-specific H10.8L shRNA design (S3 Fig). *MYH7*-R403Q hiPSC-CMs were transduced with an AAV6 viral vector expressing the H10.8L shRNA under the control of an H1 promoter. Five days post transduction, hiPSC-CMs showed a significant decrease in expression of the mutant allele RNA by allele-specific QPCR as compared to hiPSC-CMs transduced with an AAV6 vector expressing a scrambled control shRNA (Fig 2A, 57% average knockdown of mutant allele, p= 9.239e-06) as well as significant but lesser knockdown of the wild-type allele (Fig 2A 33% average knockdown of the wild-type allele, p=0.02295). These hiPSC-CMs also showed a significant decrease in their overall MYH7 levels (Fig 2B p=0.00035) while maintaining MYH6 levels comparable to untreated hiPSC-CMs (Fig 2B p=0.4121). They additionally showed significant decreases in RNA expression of hypertrophic biomarkers NPPA and NPPB (Fig 2B p= 1.736e-05 and 0.002034 respectively)

**Fig 2:**
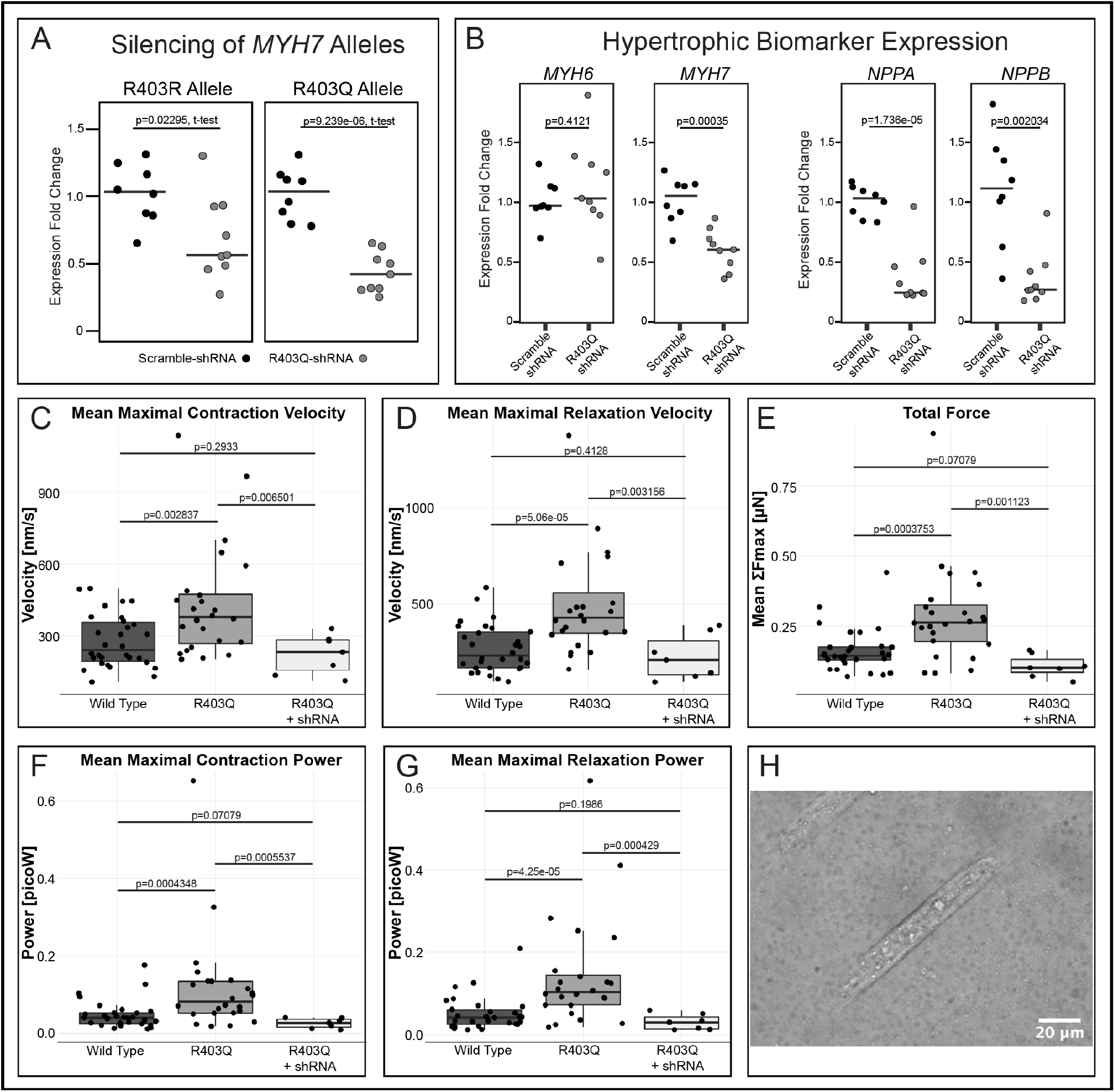
shRNA Knockdown reduces hypertrophic biomarker expression and contractile disease phenotypes in *MYH7-*R403Q hiPSC-CMs. A) Heterozygous *MYH7*-R403Q hiPSC-CMs transduced with a virally delivered R403Q-targeting shRNA (H10.8L) show reduced expression of both the R403Q and R403R alleles as compared to hiPSC-CMs transduced with a scrambled shRNA. Individual data points and medians are plotted, expression normalized to scramble-treated RNA expression for either the R403R or R403Q allele individually. hiPSC-CMs were shRNA-treated on Day 20 of differentiation, and RNA was collected after 5 days of treatment. B) *MYH7*-R403Q hiPSC-CMs treated with this same R403Q-targeting shRNA show reduction of overall *MYH7* RNA levels, but no change in *MYH6* expression levels, as well as a decrease in RNA expression of hypertrophic biomarkers *NPPB* and *NPPA*. Individual data points and medians are plotted, expression normalized to scramble-treated RNA expression. All p-values are from a two-sided t-test. C-G) Untreated *MYH7*-R403Q hiPSC-CMs from a sibling cell line (CDI Line 01178.103) show increased velocity, force, and power during contraction and relaxation compared to wild-type (WT) hiPSC-CMs, as measured by traction force microscopy. Viral transduction with the R403Q-targeting H10.8L shRNA reduce these contractile measures back to wild-type levels. H) Example image of a WT hiPSC-CM on a cellular micropatterning device. These Matrigel patterns were printed onto a polyacrylamide gel with embedded fluorescent beads to track contractile motion.

### In-vitro shRNA silencing in R403Q hiPSC-CMs shows amelioration of contractile phenotypes

*MYH7*-R403Q hiPSC-CMs from a sibling cell line derived by Cellular Dynamics Inc (CDI Line 01178.103) were cultured on protein micropatterning devices and analyzed via high-speed imagery for functional profiling using traction force microscopy. Analyses revealed that when compared to wild-type hiPSC-CMs, these *MYH7*-R403Q hiPSC-CMs had significantly increased maximal contraction and relaxation velocities and powers, as well as significantly increased maximal force (Fig 2C-G). Populations treated with the AAV6-H10.8L shRNA showed significant decreases back to wild-type levels across all measured parameters, showing an amelioration of disease phenotype.

### In-vitro ASO silencing in R403Q hiPSC-CMs shows specific knockdown of the R403Q allele

We designed allele-specific *MYH7*-R403Q-targeting antisense oligonucleotides (ASOs) based on design parameters from prior publications [30,31]. Our final ASO design comprised a 12-mer composed of 2’ O-Methyl modified RNA wings for stability, a 7-base DNA core, and a phosphorothioate backbone. When transfected into the *MYH7*-R403Q hiPSC-CMs, we saw a 49% decrease in expression of the R403Q allele (Fig 3A, p=0.001295) while we saw no significant decrease in the wild-type R403R allele as compared to hiPSC-CMs treated with a scrambled ASO. This decrease in expression was also examined using Sanger sequencing, where PCR on equal concentrations of cDNA revealed a decrease in overall expression of this region at 24 and 48 hours after sequencing with the R403Q-targeting ASO, as well as a decrease in the Sanger trace corresponding with the R403Q allele, indicating that this allele was being silenced to a greater degree than the R403R allele (S7 Fig). We also performed the same experiment examining a heterozygous SNP present on the R403Q allele and saw the same decrease in R403Q allele expression (S8 Fig).

**Fig 3:**
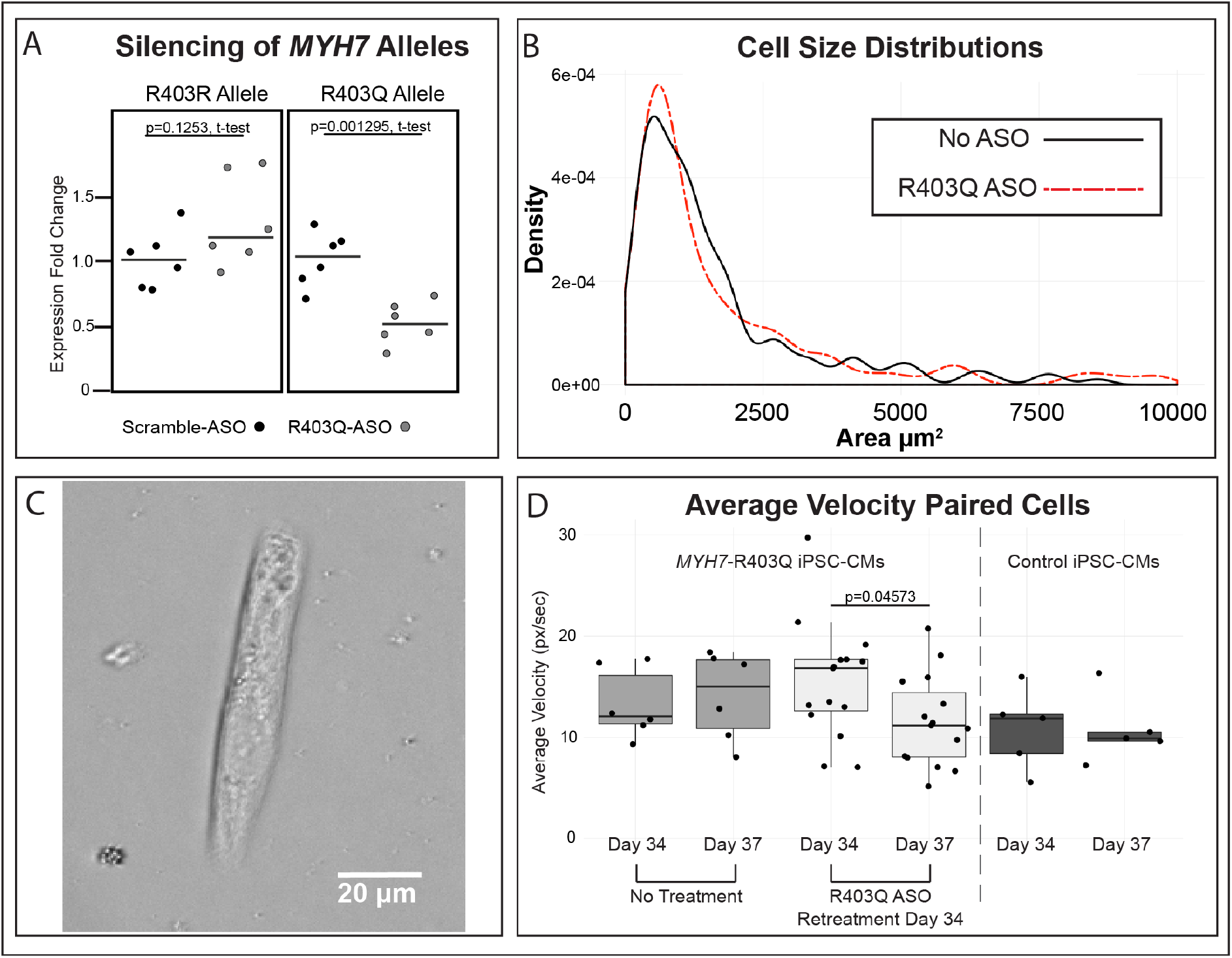
R403Q-Targeting ASO reduces contractile velocity but not cell size in *MYH7*-R403Q iPSC-CMs. A) Heterozygous *MYH7*-R403Q hiPSC-CMs transfected with an R403Q-targeting ASO show decreased expression of the R403Q allele (49% reduction, p=0.001295) but not the R403R allele. Individual data points and medians are plotted, expression normalized to scramble-treated RNA expression for either the R403R or R403Q allele individually. P-values are from a two-sided t-test. hiPSC-CMs were at differentiation day 21 at transfection, treated with 2uM ASO, and collected for RNA extraction after 24 hours. The R403Q-targeting ASO sequence is mUmCmA**CC<u>T</u>GAGG**mGmU where mN indicates 2’-O-methylated RNA bases and DNA base core is in bold. The ASO additionally has a phosphorothioate backbone. The underlined T represents the single base that discriminates the R403Q allele from the R403R allele. The scrambled control ASO used in this experiment is a 20 bp gap-mer with a fluorescent label: /56-FAM/mUmGmAmCmGTTGTACGACGmCmAmUmUmC B) Cell size density plots from the *MYH7*R403Q cell line treated with transfection-reagent only, and the *MYH7*R403Q cell line treated with the R403Q-targeting ASO during differentiation. hiPSC-CMs treated with the R403Q-targeting ASO showed a slight decrease in the median population cell size as compared to hiPSC-CMs treated with the transfection reagent alone [R403Q-ASO median = 916μm^2^, n=146 cells, vs cells treated with only transfection reagent (No ASO) median = 1077μm^2^, n=144 cells]. Plotting bandwidth adjustment = 0.8. C) Example image of an ASO-treated *MYH7*-R403Q hiPSC-CM on cellular micropatterning device used to analyze cellular motion in panel D. These Matrigel-patterns were printed onto glass-bottom dishes. The cell shown in this image is from differentiation day 34. D) *MYH7*-R403Q hiPSC-CMs were treated with the R403Q-targeting ASO at days 6, 12, and 18 of differentiation, and plated onto protein micropatterns on glass-bottom dishes on day 29. Videos taken five days later at day 34 showed that the ASOs did not have lasting effects on contraction as compared to hiPSC-CMs that received no treatment. hiPSC-CMs were retreated with the R403Q-targeting ASO after measurements were taken on Day 34, and videos were taken again 3 days later. Plotted are paired hiPSC-CMs, where we could confidently identify the same beating cell in videos taken before and after treatment. Each point represents the average contractile velocity of a single cell over a 15 second video measured in pixels per second. Untreated *MYH7*-R403Q hiPSC-CMs show a slight increase in contractile velocity over this three day time period while our ASO-retreated cell population shows a decrease in contractile velocity. Age-matched wild-type control hiPSC-CMs show little change in contractile velocity over this three day time period.

### Allele-specific ASO silencing during differentiation shows modest effects on hypertrophic cell size

Myocardial overgrowth in HCM hearts is characterized by an increase in individual cell size rather than an increase in the number of cardiomyocytes present. We aimed to reduce pathologic cell size in our *MYH7*-R403Q hiPSC-CMs via treatment with R403Q-targeting ASOs. Untreated *MYH7*-R403Q hiPSC-CMs fixed and stained at Day 30 showed an overall increase in cell population size as compared to wild-type hiPSC-CMs (S9 Fig, *MYH7*-R403Q hiPSC-CM median size = 1076μm^2^, n = 147, vs SCVI273 control cell line, median size = 808μm^2^, n = 150, p=0.002207, Wilcoxon rank-sum test without continuity correction).

We treated *MYH7*-R403Q hiPSC-CMs during differentiation, starting with a 6μM ASO treatment at day 6 of differentiation, and 3uM ASO treatments at day 12 and day 18. hiPSC-CMs were sparsely replated on day 20 and fixed at Day 30 for staining of a-actinin. hiPSC-CMs treated with the R403Q-targeting ASO showed only a modest, nonsignificant decrease in the median population cell size as compared hiPSC-CMs treated with the transfection reagent alone (Fig 3B, R403Q-ASO median = 916μm^2^, n=146 cells, vs cells treated with only transfection reagent (No ASO) median = 1077μm^2^, n=144 cells).

### Allele-Specific ASO silencing reduces contractile phenotypes of disease

To test the effects of the R403Q-targeting ASO on contractile phenotypes of disease, *MYH7*-R403Q hiPSC-CMs were treated with the silencing ASO during differentiation and plated on Matrigel micropatterns printed on glass culture dishes. Untreated *MYH7*-R403Q hiPSC-CMs showed an increase in average contractile velocity as compared to wild-type hiPSC-CMs (S10 Fig). *MYH7*-R403Q hiPSC-CMs treated during differentiation showed no relief in contractile phenotypes at Day 34 of differentiation (Fig 3D). However *MYH7*-R403Q hiPSC-CMs that were retreated with the R403Q-targeting ASO at Day 34 and imaged at Day 37 showed a decrease in average contraction velocity over the three day time period (Fig 3D). In contrast, untreated *MYH7*-R403Q hiPSC-CMs and wild-type hiPSC-CMs showed no significant change in velocity over this three day period (Fig 3D). Average contractile velocity of paired hiPSC-CMs (average ± sem): Untreated *MYH7*-R403Q hiPSC-CMs, Day 34 average: 13.3 ± 1.4 px/sec, n = 6. Untreated *MYH7*-R403Q hiPSC-CMs, Day 37 average: 14.1 ± 1.8 px/sec, n = 6. Day 34 vs Day 37 p-value = 0.7358. R403Q-ASO Treated *MYH7*-R403Q hiPSC-CMs Day 34 average: 15.5 ± 1.5 px/sec, n = 15. R403Q-ASO Retreated *MYH7*-R403Q hiPSC-CMs, Day 37 average: 11.6 ± 1.2 px/sec, n = 15. Day 34 vs Day 37 p-value = 0.04573. Wild-Type Control hiPSC-CMs, Day 34 average: 10.8 ± 1.8 px/sec, n = 5. Wild-Type Control hiPSC-CMs, Day 37 average: 10.7 ± 1.5 px/sec, n = 5. Day 34 vs Day 37 p-value = 0.9667.

## Discussion

Current therapies for hypertrophic cardiomyopathy include invasive procedures such as implantation of a defibrillator, surgical removal of heart muscle tissue (myectomy) and heart transplantation. Genetic therapeutics designed to target disease-associated alleles could present alternative, less invasive strategies of combating disease. Previous investigation of such therapeutics in HCM have focused on silencing genes in a murine model, rather than a human model. However, the dominant myosin isoforms are different between mice and humans. Here, we present the examination of two potential genetic therapeutics targeting a heterozygous disease-causing mutation in *MYH7* as tested in a human iPSC-CM model of disease, using biophysical assays to assess their impacts on contractile phenotypes of disease.

To assess allele-specific silencing via shRNA of an HCM-associated gene in human cells, we introduced an shRNA designed to specifically target the *MYH7*-R403Q allele into hiPSC-CMs from a patient heterozygous for this mutation. Previous studies using patient-derived hiPSC-CMs have shown them to be a useful model for studying disease-phenotypes in the laboratory and have proven to be an important platform for studying potential silencing therapeutics in cardiovascular disease [18,20]. The patient from whom our cell line was derived was diagnosed with nonobstructive HCM at age 13 and was recommended to have an ICD placed at age 39 due to nonsustained ventricular tachycardia. This patient additionally had a strong family history of early death due to heart failure from HCM as well as family members who additionally carried the R403Q mutation and one who had undergone a heart transplant.

After viral transduction with the R403Q-targeting shRNA, hiPSC-CMs containing the heterozygous *MYH7*-R403Q mutation showed 57% knockdown of the mutant allele as measured by allele-specific qPCR. We additionally saw a decrease in molecular markers of hypertrophy, including *NPPA* and *NPPB*. Previous studies of allele-specific *MYH6* silencing in mice have shown similar decreases in *NPPA* and *NPPB* expression, further supporting the idea that reducing mutant myosin transcripts may have beneficial effects on other cardiomyopathy-related gene expression [15]. While we do see a 41% reduction of overall *MYH7*, we do not see a reduction in *MYH6*, indicating that our shRNA treatment is specific to *MYH7* and does not cause off-target silencing of *MYH6*.

To assess functional improvement in phenotype of shRNA-treated hiPSC-CMs, we assayed contractile function of a related *MYH7*-R403Q hiPSC line. We deployed cellular micropatterning devices on hydrogel substrates to measure contractile properties of hiPSC-CMs using traction force microscopy (as described in Ribeiro, et al. [28]). Previous studies of hiPSC-CMs and cardiac tissues derived from patients with HCM have shown contractile dysfunction, including contractile arrhythmias and hypercontractility [20,32,33], as well as greater developed force and shorter twitch duration [34], in patient-derived cardiomyocytes as compared to wild-type controls. In our study, we found that *MYH7*-R403Q hiPSC-CMs had significantly increased maximal contraction and relaxation velocities and powers, as well as significantly increased maximal force when compared to wild-type hiPSC-CMs. We saw a shift towards wild-type hiPSC-CM levels of all measured parameters in *MYH7*-R403Q hiPSC-CMs after shRNA treatment, as compared to non-transfected *MYH7*-R403Q hiPSC-CMs. While our shRNAs reduced expression of not only the mutant (R403Q) but also the wild-type (R403R) *MYH7* allele in our in-vitro studies, this overall drop in *MYH7* did not prevent the return to normal phenotype in our silenced hiPSC-CMs, leading us to believe that the small drop in *MYH7*-R403R RNA is tolerated.

We continued to pursue an allele-specific silencing molecule, and turned to antisense oligonucleotides, which have shown great promise in both neurodegenerative disorders and cardiovascular disease. We tested a number of gap-mer ASO designs, from 12-20 bp in length, with the target variant base either centered or slightly-off center, and with or without additional mismatches, before settling on our final 12 bp design which specifically silenced the R403Q allele (S5 Fig & S6 Fig). Our best designed ASO showed a 49% decrease in *MYH7*-R403Q expression with no significant knockdown of the wild-type allele (Fig 3A). However, while this short ASO design lended itself to the greatest specificity, it does pose potential problems when considering off-target silencing. A recent paper by Yoshida et al. estimated that 12-bp ASOs may have up to 3.4 potential off-target binding regions with zero-sequence mismatches, and up to 122 potential off-target binding regions if a single mismatched-base is tolerated [35]. While these numbers are large, they do not take into account potential secondary structure of the RNA, which is known to impact target-affinity in ASOs [36], or whether or not the transcripts containing those potential off-target sequences are expressed in the target cell-type. Further investigation of the effects of these ASOs on off-target gene expression via whole transcriptome RNA-expression analysis at both the cell and tissue levels would be necessary before this oligonucleotide was introduced into a clinical setting.

Despite these considerations, we aimed to reduce pathologic cell size in our *MYH7*-R403Q hiPSC-CMs via treatment with the R403Q-targeting ASOs. We showed that Day 30 *MYH7*-R403Q hiPSC-CMs showed an increase in overall population cell size as compared to a wild-type line (S9 Fig). We found that treatment of *MYH7*-R403Q hiPSC-CMs during differentiation with the R403Q-targeting ASO could induce a modest reduction in overall population cell size at Day 30 (Fig 3B), as compared to hiPSC-CMs treated with only the transfection reagent. Previous examinations of cell size in hypertrophied human hearts have shown mean cell diameter changes of around 25% in the left ventricular walls as compared to control hearts [37]. Our untreated *MYH7*-R403Q hiPSC-CMs show a 33% increase in median cell size as compared to the wild-type control line. However our R403Q-ASO treated hiPSC-CMs show only a 13% increase in median cell size (Fig 3B), showing modest relief of phenotype back towards wild-type cell sizes.

As with our shRNA treated hiPSC-CMs, we aimed to examine the effects of our antisense oligonucleotide silencing on contractile phenotypes in our hiPSC-CMs. We used automated video analysis to assess the impact of the R403Q-targeting ASO on contractile phenotypes of disease. While ASO treatment during differentiation did not have lasting effects on contraction, this was not especially surprising, given that we see persistence of ASO silencing in our hiPSC-CMs for around one week, and the hiPSC-CMs had not received a dose of ASO for 16 days by the analysis time point. However when we retreated hiPSC-CMs at this time point and imaged again three days later, we saw a decrease in the average contractile velocity of hiPSC-CMs treated with our allele-specific ASO (Fig 3D). We did not see a decrease in the velocity of untreated *MYH7*-R403Q hiPSC-CMs over the same 3 day period (Fig 3D). While we examined paired hiPSC-CMs in Fig 3D, where we were able to confidently match videos of hiPSC-CMs taken at Day 34 and Day 37, we also present population level data of all beating hiPSC-CMs recorded for these populations in S10 Fig. This may suggest that ASO silencing is able to relieve contractile phenotypes after cell growth and maturation.

A growing number of investigations of HCM-associated *MYH7* mutations have also studied the allele-specific expression of both wild-type and mutant *MYH7* alleles, and have reported imbalanced expression not only between patients, but between individual cells from the same patient [38–40]. These imbalances have been associated with variable contractile and calcium-handling phenotypes between mutation-containing cardiomyocytes [38], with some cells showing properties quite similar to wild-type cells and others showing highly disordered phenotypes. Montag et al. suggests that these imbalances may be due to cells with high wild-type-allele expression mimicking healthy cells and cells with high mutant-allele expression exhibiting more detrimental phenotypes [38]. This underlying imbalance in expression between cells may not only contribute to the increased difficulty in phenotyping large cell populations, where variance between cells may make population level conclusions noisy, but also to the disconnect between allele-specificity and phenotyping effect that we observe in our silenced hiPSC-CM populations.

While the R403Q-targeting shRNA knocked down both alleles of *MYH7*, it conferred significant benefits to both molecular and contractile phenotypes of disease. Yet while the R403Q-targeting ASO showed more specific knockdown, it also showed more modest phenotypic benefits. This disparity in expected phenotypic response may be due to the dose and delivery of each molecule. AAV viral delivery results in episomal stability of the delivered genetic material, allowing high levels of the shRNA to be transcribed in the cell under the control of the H1 promoter. Conversely, despite delivering our ASOs at high concentrations (6μM) we found their silencing effects to last for only about one week (S7 Fig). It may be that persistent treatment with an *MYH7* RNA-targeting molecule is needed in order to sufficiently silence the disease-associated transcripts. This would pose important considerations for future clinical applications, where repeated treatments can become expensive and burdensome to patients and caretakers. Additionally, our R403Q-targeting ASO decreased mutant allele expression by only about half, leaving dominant-negative transcripts present in the cell. While we created a number of other ASOs during the design process that knocked down overall *MYH7* to much greater levels, none of them proved to be allele-specific (S5 Fig & S6 Fig). This tradeoff between specificity and overall silencing may be important to consider in situations where even a small amount of mutant protein may be enough to cause disease. Single, efficient treatment with an shRNA based strategy, as has been done in related animal models of hypertrophic cardiomyopathy [16] may prove to be the most efficacious path forward.

In conclusion, therapeutic gene silencing has the potential to ameliorate disease phenotypes by targeting the underlying genetic cause of disease. Here, we examined two methods of gene silencing in a human iPSC-CM disease model of hypertrophic cardiomyopathy, shRNA and ASO silencing. We used traction force microscopy and automated video analysis to interrogate the effects of these silencing methods on contractile disease phenotypes. We found dissociation of allelic specificity and functional improvements, suggesting a more complex allelic control underlying the role of *MYH7*-R403Q in disease expressivity. We demonstrated that while less allele-specific, shRNA silencing can significantly improve contractile and molecular phenotypes of disease, potentially due to high, continuous levels of treatment conferred by AAV viral delivery. We additionally showed that while ASO silencing has the potential to improve silencing allele-specificity, it showed more mild relief of phenotypes in our hiPSC-CMs. These findings demonstrate that decreasing expression of *MYH7* in a human model of disease has the potential to relieve hypertrophic cardiomyopathy phenotypes, but that further study may be needed to parse the relationship between therapeutic silencing and mutant and wild-type expression underlying disease phenotypes.

## Materials and methods

### *MYH7*-R403Q induced pluripotent stem cell derivation and culture

hiPSCs were derived from two siblings with the *MYH7*-R403Q mutation. hiPSCs for the first line were derived from patient fibroblasts collected under Stanford Institutional Review Board GAP (Genetics and Proteomics) approval number 4237 and cultured under Stanford SCRO (Stem Cell Research Oversight) 568 (Fig 1A, patient 1). Additional experiments were carried out under Stanford Protocol 2663. Derivation of patient-specific hiPSC lines was performed as previously described [20,41]. Fibroblasts were passaged before infection with lentiviral vectors containing the four Yamanaka factors (OCT4, SOX2, KLF4, and c-MYC). hiPSC colonies were maintained on Matrigel-coated plates (BD Biosciences) in mTESR-1 medium (StemCell Technologies) [20]. These cells were used for all silencing experiments except Fig 2C-G. Experiments in Fig 2C-G were conducted on a second line from a sibling (Fig 1A, patient 2) who also carried the *MYH7*-R403Q mutation. hiPSCs and hiPSC-CMs for this line were derived by Cellular Dynamics International (Line 01178.103). hiPSCs were maintained on 1:200 Matrigel plates in mTesr. The wild type control cell line used for all ASO silencing experiments was line SCVI273 obtained from the Stanford Cardiovascular Institute Biobank.

### hiPSC-cardiomyocyte differentiation

hiPSCs were differentiated to cardiomyocytes using a modified version of Sharma et al. 2015 [42]. Briefly, hiPSCs were cultured until they reached 80% confluence in 12-well dishes. For ASO phenotyping experiments: hiPSCs were then exposed to differentiation media (RPMI with B27 - Insulin) for two days with the addition of GSK3 inhibitor CHIR. Optimal CHIR concentrations varied between cell lines. Each new cell line was tested with concentrations ranging from 3μM to 8μM. Cells were then exposed to differentiation media with the addition of Wnt inhibitor IWR for two days. After that, cells were cultured in the media alone for two days before changing to RPMI with complete B27 for 3 days. hiPSC-CMs were then maintained in starvation media (RPMI minus glucose with B27 and lactate added). This helped to purify and mature only the hiPSC-CMs which can metabolize lactate in the absence of glucose. hiPSC-CMs were cryopreserved in BamBanker and stored in liquid nitrogen. For shRNA experiments and Fig 3A, differentiation differed slightly with a single day “rest” period between CHIR and IWR exposure, and with maintenance of hiPSC-CMs in media containing glucose after Day 20.

### hiPSC-cardiomyocyte shRNA silencing experiments

hiPSC-CMs were seeded into a 24-well plate at a density of 500K cells per well. hiPSC-CMs were transduced at differentiation day 20 in 300μl of RPMI/B27 media using 3.9e^9^vg of AAV6-scramble-shRNA or AAV6-H10.8L shRNA expressing virus (S2 Fig). Nine wells were transfected with the scramble-shRNA virus while ten wells were transfected with the H10.8L virus. hiPSC-CMs were collected after 5 days and RNA was extracted with the Qiagen miRNeasy kit, with additional DNase treatment step. cDNA was made with the Applied Biosystems High Capacity cDNA kit. QPCR was performed as described below. One sample from each condition in Fig 1A-B was excluded due to high outlier values in *MYH7* QPCR and abnormal housekeeping gene QPCR values. The H10.8L-containing AAV6 virus was produced at Stanford’s Neuroscience Gene Vector and Virus Core while the scramble-shRNA-containing AAV6 virus was produced by Virovek, Hayward, CA.

### QPCR analysis of MYH7 alleles

Allele specific QPCR was performed using mutant or wildtype specific forward primers and a common reverse primer, as previously described in Wheeler et al. [41]. Each forward primer contains a mismatch at the penultimate nucleotide to increase allele specificity. The *MYH7* specific fluorescent probe was optimized for maximum sequence dissimilarity from *MYH6*. Allele specific qPCR conditions using Taqman Fast Universal PCR Master Mix: 95°C 20”, 40 cycles of 95°C 30”, 58°C 20”, 72°C 30” for R403Q or 40 cycles of 95°C 30”, 64°C 20”, 72°C 30” for R403R. Endogenous control was EEF1A2. Data was analyzed using the delta-delta Ct method.

R403R Forward Primer: 5’ GGGCTGTGCCACCCTAA 3’
R403Q Forward Primer: 5’GGGCTGTGCCACCCTAG 3’
Common Reverse Primer: 5’CGCGTCACCATCCAGTTGAAC 3’.
*MYH7* specific fluorescent probe: FAM-5’TGCCACTGGGGCACTGGCCAAGGCAGTG 3’-TAMRA.

### RNA extraction and cDNA amplification

For some early experiments, RNA was extracted using the Qiagen miRNeasy Kit with on-column DNAse treatment. However for the majority of experiments, RNA was extracted using trizol/chloroform and treated with the Thermo Fisher Turbo DNAse kit. cDNA amplification was achieved using the High Capacity cDNA Reverse Transcription Kit with RNase Inhibitor. For some reactions, half of the cDNA was digested with AvaI (to specifically cut the wild-type allele) and the other half was cut with Bsu36I (to specifically cut the mutant allele) before their individual QPCR reactions, though we later determined this step to be nonessential.

### QPCR of hypertrophic genes and biomarkers

QPCR of common hypertrophic and biomarker genes was performed using IDT predesigned probes and IDT PrimerTime Master Mix on a Viia7 thermocycler. Individual probes are listed in Table S2. Data was analyzed using the delta-delta Ct method.

### Micropatterning of single cells on polyacrylamide gel surfaces and analysis of contractility from traction force microscopy

Micropatterning extracellular proteins on a surface for cell culture can set the morphology of single cells to match the same shape of the micropatterned region. Culturing single hiPSC-CMs on micropatterns with a rectangular shape forces these cells to have a more mature morphology and organization of the sarcomere-based contractile machinery [28]. In addition, micropatterning hiPSC-cardiomyocytes with a rectangular shape on a soft material with physiological tissue rigidity further enhances the physiology of their contractile function and allows measurement of contractility by tracking the cell-induced deformation of the material under each cell [27,28]. *MYH7*-R403Q hiPSC-cardiomyocytes from CDI Line 01178.103 and healthy control line 01027.101 were cultured on 2000 μm^2^ rectangular (7:1 aspect ratio) Matrigel patterns on polyacrylamide (PA) surfaces as previously described in Ribeiro et al. [28,41,43]. Soft lithography was used to fabricate PDMS microstamps [44]. Microstamps were flooded with 1:10 dilution of Matrigel in cold L15 medium at 4°C for 24 hours and dried using N_2_ after aspirating and washing the surface twice with cold L15 medium. Patterns were stamped onto a plasma-treated glass coverslip. Gels were fabricated as reported elsewhere [41,45,46]. PA gel components (12% acrylamide, 0.15% N, N-methylene-bis-acrylamide) and fluorescent microbeads were mixed in DI water. 50 μl of the solution was added to clean, pretreated coverslips. Ammonium persulfate was used as a catalyst for gel polymerization and TEMED was used as an initiator. Micropatterned coverslips were then placed on top of the gel solution. After polymerization, coverslips were removed to reveal the patterns, and hiPSC-CMs were seeded onto them in appropriate culture media. *MYH7*-R403Q hiPSC-CMs were transduced with 3.9e^10^vg of AAV6-scramble-shRNA or left untreated. Videos of beating hiPSC-CMs were acquired with a Zeiss Axiovert 200M inverted microscope equipped with a Zeiss Axiocam MRm CCD camera to image the displacement of fluorescent microbeads in polyacrylamide materials supporting the adhesion of micropatterned cells. The microscope had an environmental chamber (PeCon) to keep temperature of cell media at 37 °C. Videos of beating hiPSC-CMs were acquired while electrically pacing the hiPSC-CMs with 10 ms-wide bipolar pulses of electric-field stimulation at 10-15 V with a frequency of 1 Hz or 2 Hz (Myopacer, IonOptix). Only cells beating at the appropriate frequency were analyzed. We measured the parameters of cell contractility from traction force microscopy results following the methodology reported by Ribeiro et al [47].

### hiPSC-CM staining for cell-size measurements

hiPSC-CMs were stained for α-actinin (to confirm cardiomyocyte identity), actin, and DAPI for cell-size measurements. Briefly, beating hiPSC-CMs plated on Matrigel-coated coverslips were relaxed in 50mM KCl before fixation with 4% paraformaldehyde. hiPSC-CMs were permeabilized with 0.1% Triton X-100 and incubated in 5% normal goat serum before overnight incubation with the primary α-actinin antibody (Sigma Aldrich, A7811) at 4C. After washing with 0.1% Tween-20, hiPSC-CMs were incubated for one hour with the secondary antibody (Alexa Fluor 568, Sigma Aldrich, A11031). After washing again with Tween-20, hiPSC-CMs were incubated in ActinGreen for ten minutes before mounting with Prolong Diamond Antifade Mountant with DAPI. The next day, hiPSC-CMs were imaged using a Nikon Eclipse 90i microscope. Cell-size measurements were calculated by blinded circling of hiPSC-CMs in ImageJ. Cell sizes greater than 10,000μm^2^ were removed prior to analysis.

### ASO silencing in hiPSC-cardiomyocytes

ASO silencing experiments used different ASOs, concentrations, and transfection reagents as described in the text. However, the base protocol is as follows. ASOs were custom-designed and ordered from IDT (Integrated DNA Technologies, Coralville, IA) before resuspension to 100μM in sterile H_2_O. hiPSC-CMs or differentiating hiPSCs were plated in 12 or 24-well tissue culture plates on 1:200 Matrigel. After 2-3 days, culture media was removed and replaced with fresh media. For each ASO, a master mix was prepared. The transfection reagent (TransIT-TKO, Mirus Bio LLC) was added to fresh Opti-MEM and mixed gently. The ASO was added to the desired concentration and mixed gently by pipetting before sitting at room temperature for 30 minutes. As an example, for 1 well of 6μM ASO in a 12-well plate, 49μl of Opti-MEM was mixed with 3μl of TransIT-TKO before the addition of 48μl of ASO. After 30 minutes, 100μl of the master mix was added dropwise to the appropriate well of the tissue culture dish, which contained 700μl of fresh media. The culture plate was gently shaken to disperse the ASO complexes and returned to 37C for at least 24 hours. When collected for RNA analysis, hiPSC-CMs were either dissociated with accutase, rinsed in PBS, and centrifuged or collected with trizol. Pellets or trizol collections were snap frozen in liquid nitrogen before RNA extraction. ASOs were ordered from Integrated DNA Technologies, and have a phosphorothioate backbone. R403Q-targeting ASO sequence: mUmCmA**CCTGAGG**mGmU where mN indicates 2’-O-methylated RNA bases and DNA base core is in bold. Scramble ASO sequence used in Fig 3A, designed using InvivoGen’s siRNA wizard: /56-FAM/mUmGmAmCmGTTGTACGACGmCmAmUmUmC.

### Silencing through differentiation treatments

ASO silencing treatments during differentiation were performed as described above, except that treatments were delivered throughout differentiation at days 6, 12, and 18 in 12-well dishes. The day 6 treatment consisted of 6μM ASO and 3μl of TransIT-TKO while the day 12 and day 18 treatments were halved (3μM ASO and 1.5μl TransIT-TKO per well) to avoid toxicity due to repeated high exposure to transfection reagent.

### Cellular micropatterning on glass for velocity analysis

Microcontact printing of rectangular protein patterns was performed as previously described [48]. Briefly, PDMS microstamps containing 17×119 mm^2^ rectangles were used to microprint dilute Matrigel onto glass bottom 6-well plates. hiPSC-CMs were replated onto micropatterns at 10,000-50,000 cells/mL. Videos of patterned hiPSC-CMs were acquired using a Keyence BZ-X microscope. Videos were taken using a 40X objective for 15 seconds at 29 frames per second. Videos were analyzed as described in Huebsch et al [29]. Average maximum velocity was calculated for each cell. For each cell with an average maximum velocity above 30 pixels/second, a secondary round of examination was performed to remove any cell suspected of being two hiPSC-CMs attached to the same pattern. These doublet hiPSC-CMs were removed from all further analysis.

### Statistical analysis

All statistical analysis was performed using RStudio. QPCR gene and allele expression data was analyzed using Student’s t-test. Contractility measurements in Fig 2C-G were analyzed using a two-tailed Mann-Whitney U test. Cell size measurements in Fig 3B were analyzed using a Wilcoxon rank-sum test without continuity correction. Velocity measurements in Fig 3D were analyzed using Student’s two-sided t-test.

## Notes

Financial Support A. Dainis received support from the NSF Graduate Research Fellowship Program. E. Ashley received funding from NIH Director’s New Innovator Award DP2 OD004613 and is supported by NIH U24 award 1U24EB023674-01 and NIH U01 award 1U01HG007708. B. Pruitt received support from AHA 1205987-120-UAKOD as well as NIH 1R21HL13099301. A. Ribeiro received support from AHA Fellowship 14POST18360018. J.C. Wu is supported by NIH R01 HL130020 and NIH R01 HL126527. The funders had no role in study design, data collection and analysis, decision to publish, or preparation of the manuscript.

Disclosures E. Ashley is a Founder of Personalis and DeepCell, Inc, and an advisor for SequenceBio. M. Wheeler is an equity partner in Personalis, Inc and has been a consultant to MyoKardia.

