## Supplemental Material for "Dissociation of disease phenotype and allele silencing in hypertrophic cardiomyopathy"

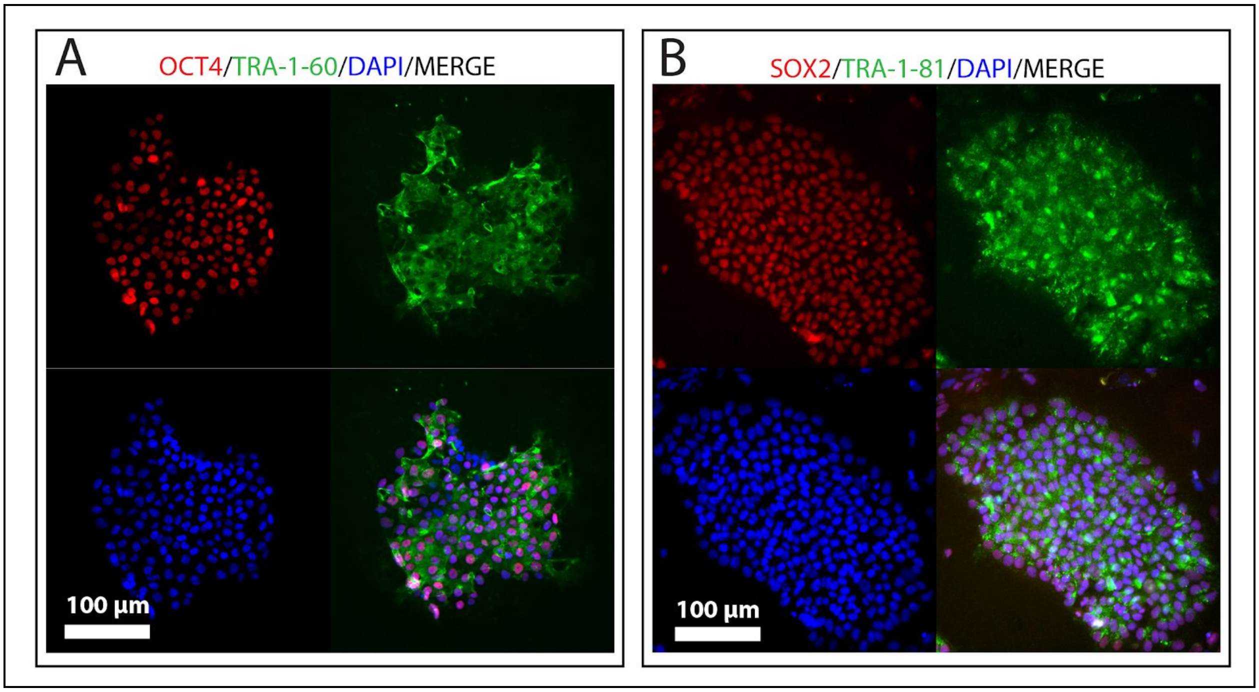


**S1 Fig: Pluripotency marker staining in *MYH7*-R403Q hiPSCs:** Immunofluorescent staining of pluripotency markers in hiPSCs derived from a patient containing the *MYH7*-R403Q mutation. Panel A represents staining of OCT4 and TRA-1-60 while panel B shows staining of SOX2 and TRA-1-81. Expression of these markers indicated successful derivation of hiPSCs from fibroblasts.

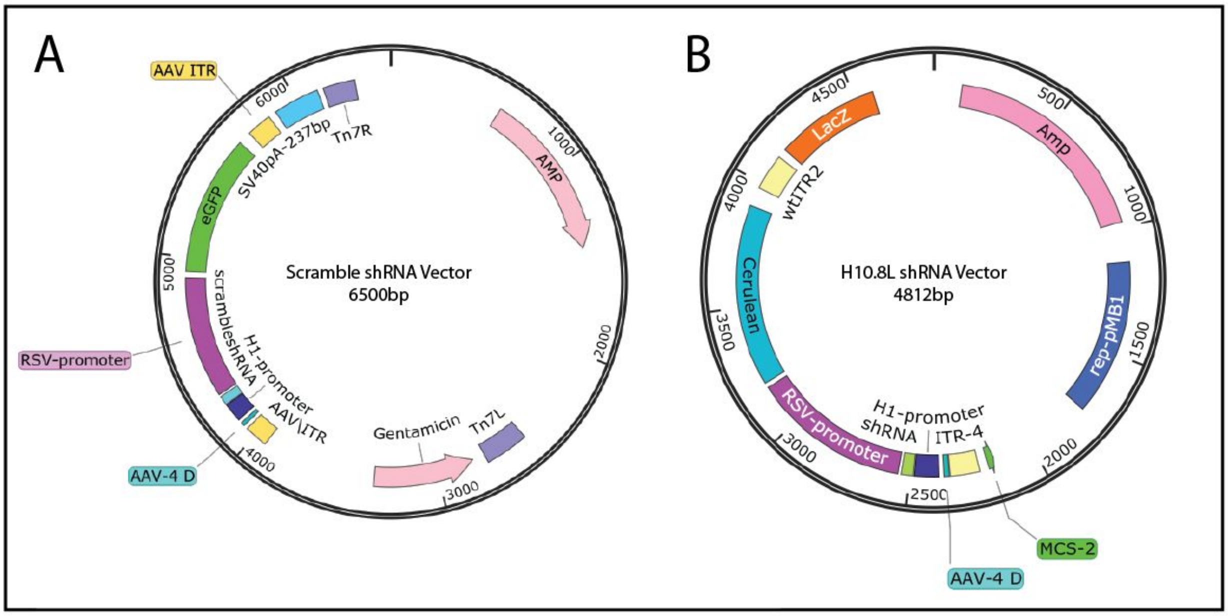


**S2 Fig: Viral vectors containing scramble and H10.8L shRNAs** Viral vectors containing either a scrambled shRNA or the H10.8L R403Q-targeting shRNA were used to create AAV6 viral particles for transduction. Each shRNA is under the control of the H1-promoter, and each vector contains a fluorescent marker under the control of an RSV promoter (eGFP for the scramble shRNA vector and Cerulean for the H10.8L vector). These cassettes resided between AAV inverted terminal repeats (ITR), critical to multiplication of the viral genome, with additional selective markers outside of the ITRs for molecular biology and cloning purposes. The scramble shRNA sequence is: GGCCAGCCTGCGTATCATAgaagcttgTATGATACGCAGGCTGGCC where the lowercase section represents the hairpin loop.

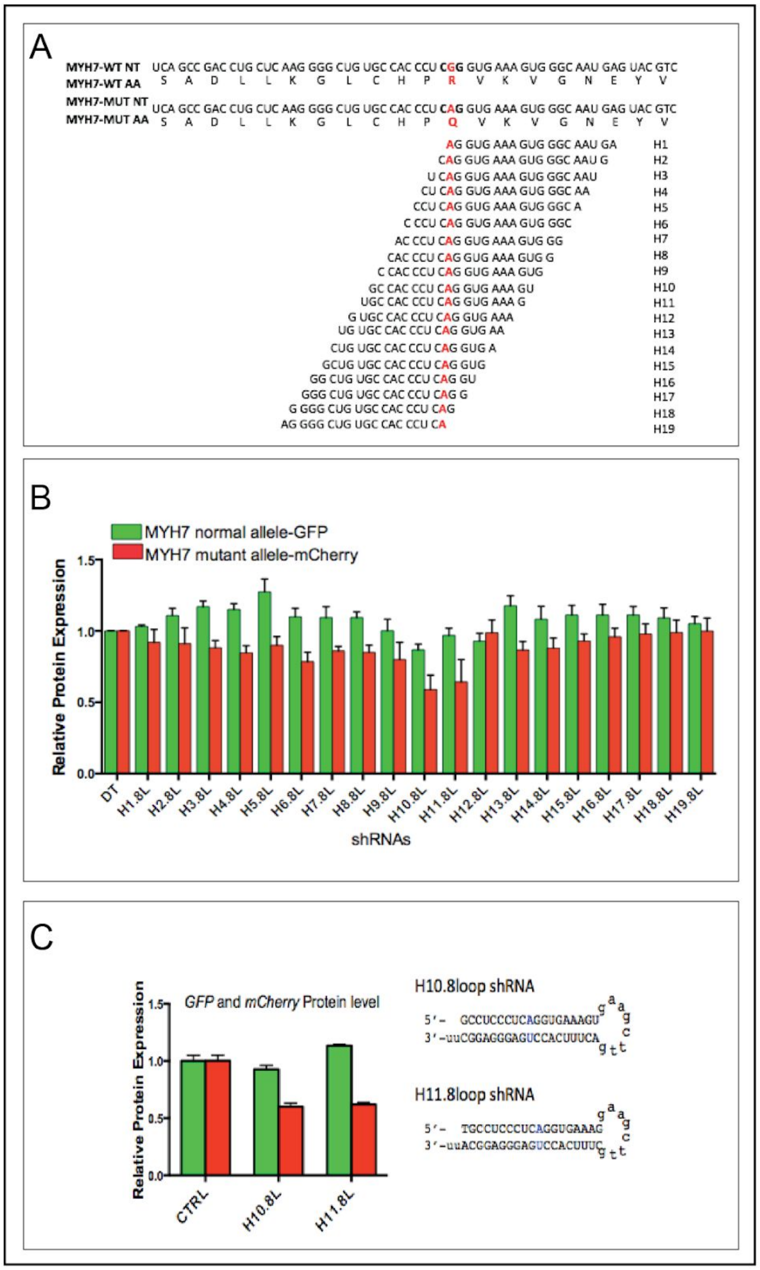


**S3 Fig: Complete siRNA "walk" design targeting *MYH7*-R403Q**

A) siRNA walk design: List of siRNAs designed to target the R403Q allele in vitro, aligned to the mutant allele, to demonstrate the one-base displacement between each siRNA. B) HEK cells were transfected with two plasmids: one containing a truncated *MYH7*-R403R transcript fused to GFP and one containing a truncated *MYH7*-R403Q transcript fused to mCherry. The doubly-transfected HEK cell population was then transfected with shRNAs containing each of the sequences shown in panel A connected by an 8 base pair loop to form a hairpin. Plotted are Fluorescence Activated Cell Sorting (FACS) quantification of GFP and mCherry in each shRNA treated cell population. C) Protein quantification of GFP and mCherry reporters using Fluorescence activated cell sorting FACS after transfection with the two shRNAs that showed the highest allele-specific discrimination in panel B, H10.8L and H11.8L shRNAs. The full hairpin design of H10.8L and H11.8L shRNAs are also shown. Data in this figure previously produced in US Patent Application US20160348103A1 [1].

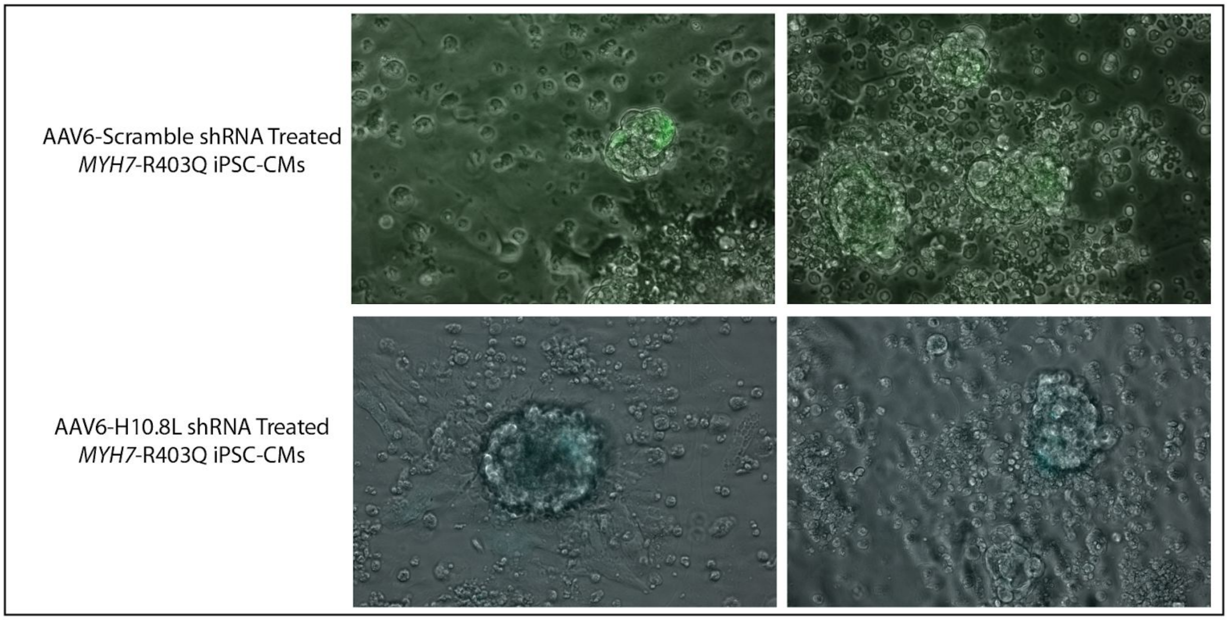


**S4 Fig: Representative images of *MYH7*-R403Q hiPSC-CMs treated with either the scramble shRNA or H10.8L shRNA virus**

Representative images of *MYH7*-R403Q hiPSC-CMs transduced with either the AAV6 scramble-shRNA virus, which contained an eGFP cassette, or the AAV6 H10.8L (R403Q-targeting) shRNA virus, which contained a Cerulean cassette. Images were taken 5 days after transduction, 1 hour before collection for RNA extraction in Fig 2A. Images have been autoscaled for brightness.

| ASO | Design |
|---|---|
| Scramble-F ASO | /56-FAM/*mUmGmAmCmG*TTGTACGACG*mCmAmUmUmC* |
| R403Q 20mer ASO | *mAmCmUmUmU*CACCTGAGGG*mUmGmGmCmA* |
| R403Q_Mismatch1 | *mAmCmUmUmU*CAC**G**TGAGGG*mUmGmGmCmA* |
| R403Q_Mismatch2 | *mAmCmUmUmU*CACCT**C**AGGG*mUmGmGmCmA* |
| R403Q_Mismatch3 | *mAmCmUmUmU*CA**G**CTGAGGG*mUmGmGmCmA* |
| R403Q_Mismatch4 | *mAmCmUmUmU*CACCTG**T**GGG*mUmGmGmCmA* |
| R403Q_Shift1 | *mCmUmUmUmC*ACCTGAGGGT*mGmGmCmAmC* |
| R403Q_Shift2 | *mCmAmCmUmU*TCACCTGAGG*mGmUmGmGmC* |
| R403Q_Shift3 | *mUmUmCmAmC*CTGAGGGTGG*mCmAmCmAmC* |
| R403Q_Shift4 | *mCmCmCmAmC*TTTCACCTGA*mGmGmGmUmG* |
| R403Q_12mer_7gap | mU*mC*mA*C*C*T*G*A*G*G*mG*mU |

**S1 Table: Sequences of additionally tested antisense oligonucleotides**

In our efforts to create an allele-specific ASO, we experimented with a number of different ASO designs. These included designs that incorporated mismatches to decrease binding affinity to off target locations as well as shifting the ASO's placement relative to the target base position. To begin, we designed a 20mer ASO targeting the R403Q allele (denoted as R403Q 20mer ASO). All of our designed ASOs used a "gap-mer" structure which consists of a DNA core and modified RNA "wings" for stability. These designs were based on prior promising allele-specific silencing results from the literature [2–4]. We found that exposing our *MYH7*-R403Q hiPSC-CMs to this R403Q 20mer ASO at 2uM concentrations resulted in an almost total decrease in expression of both the R403Q and R403R alleles [S5 Fig]. We then designed a number of ASOs that used the R403Q ASO as a starting design, but included single base pair mismatches to decrease binding affinity to off target loci. Stadhouders et al. reported that A-A, A-G, C-C, G-G, G-A binding mismatches showed the greatest decrease in off target binding in a real-time PCR assay, so we designed our mismatch ASOs to contain these mismatch pairs when possible [5]. We found that none of these ASOs showed allele-specific silencing of the R403Q allele [S5 Fig]. We also tested a number of ASOs without mismatches but that were shifted 1 or 3 bases to the left or right relative to the target SNP, while maintaining a 20mer length. We again found that none of these ASOs showed allele-specific silencing of the R403Q allele [S5 Fig]. We additionally modified ASOs to contain a phosphorothioate backbone, which has been previously shown to increase both ASO stability and uptake into cells [6]. We created a phosphorothioate backbone ASO 12bp in length. Our 12mer ASO showed specific silencing of the R403Q allele ["R403Q 12mer 7gap", S6 Fig], and is the R403Q-targeting ASO used in the main body of the manuscript. Listed here are the sequences of ASOs designed with "mismatches," "shifts," and differing lengths in an attempt to improve allele-specificity. Italicised sections indicate 2' O-Methyl modified RNA wings. The target SNP (R403Q) is the underlined "T" in these sequences. The "mismatches" designed to decrease off-target binding affinity are indicated in bold. The * represents a phosphorothioate backbone.

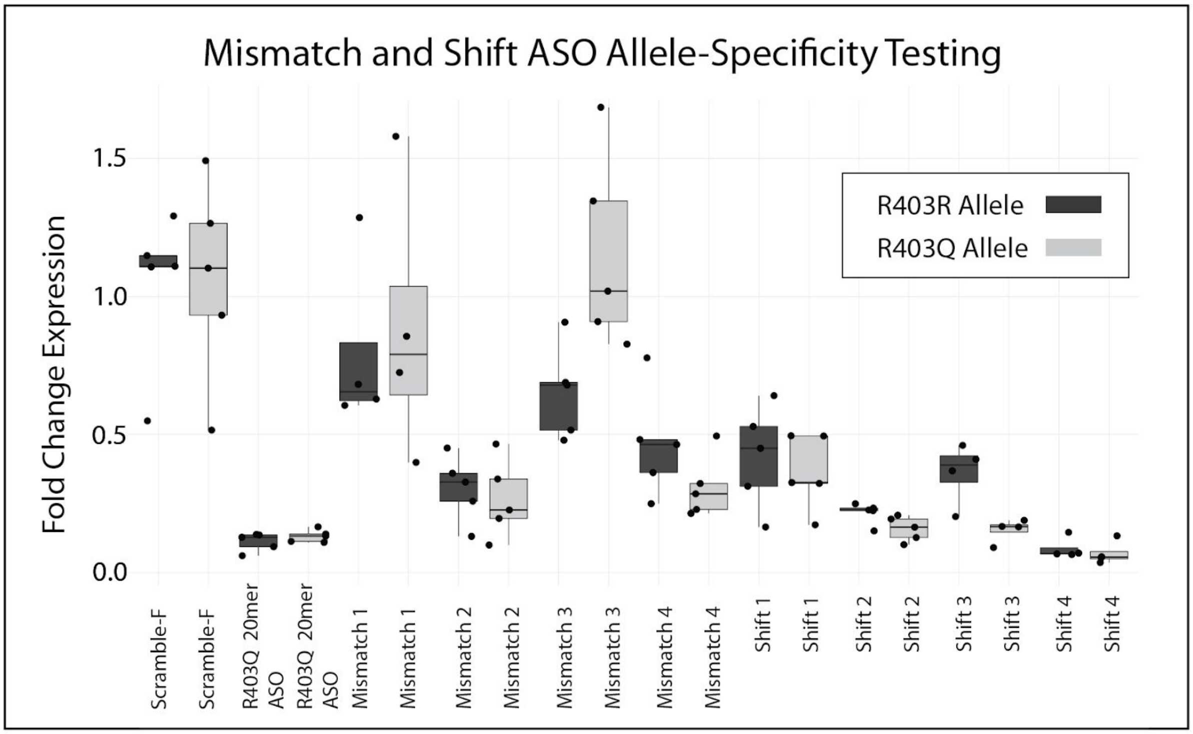


**S5 Fig: Allele-specificity testing of mismatch and shift ASOs** Allele specific expression of the wild-type R403R or mutant R403Q alleles in *MYH7*-R403Q hiPSC-CMs treated with our range of mismatch and shift ASOs from S1 Table. Expression of each allele is individually normalized to expression of that allele (either R403R or R403Q) in cells treated with a scrambled ASO. Each point represents an individual well of treated cells. We do not see R403Q-specific silencing in any of our treated populations, though we do see a range of overall expression levels.

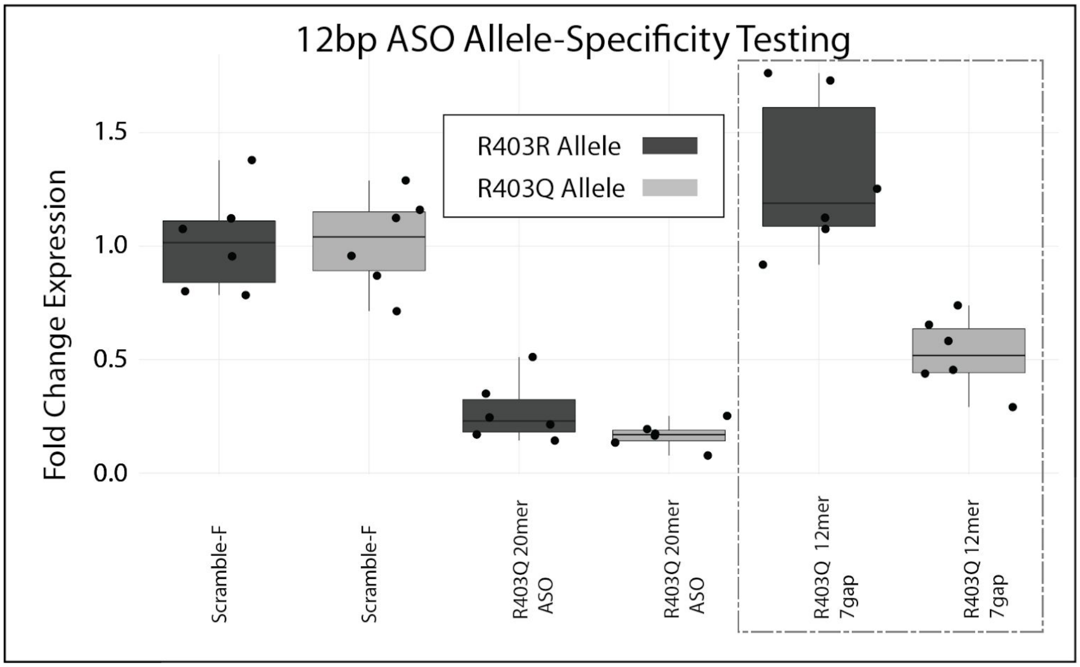


**S6 Fig: Allele-specific silencing with phosphorothioate backbone ASO** Allele specific expression of the wild-type R403R or mutant R403Q alleles in *MYH7*-R403Q hiPSC-CMs treated with our range of mismatch and shift ASOs from S1 Table. Expression of each allele is individually normalized to expression of that allele (either R403R or R403Q) in cells treated with a scrambled ASO. Each point represents an individual well of treated cells. We see allele specific-silencing of the R403Q allele in cells treated using the R403Q 12mer 7gap ASO (an ASO with a total length of 12 bp with a 7 bp DNA core). This ASO shows no silencing of the R403R allele, and became our R403Q-targeting ASO used in the main text. This data, without the 20mer ASO for comparison, is the data that appears in Fig 3A.

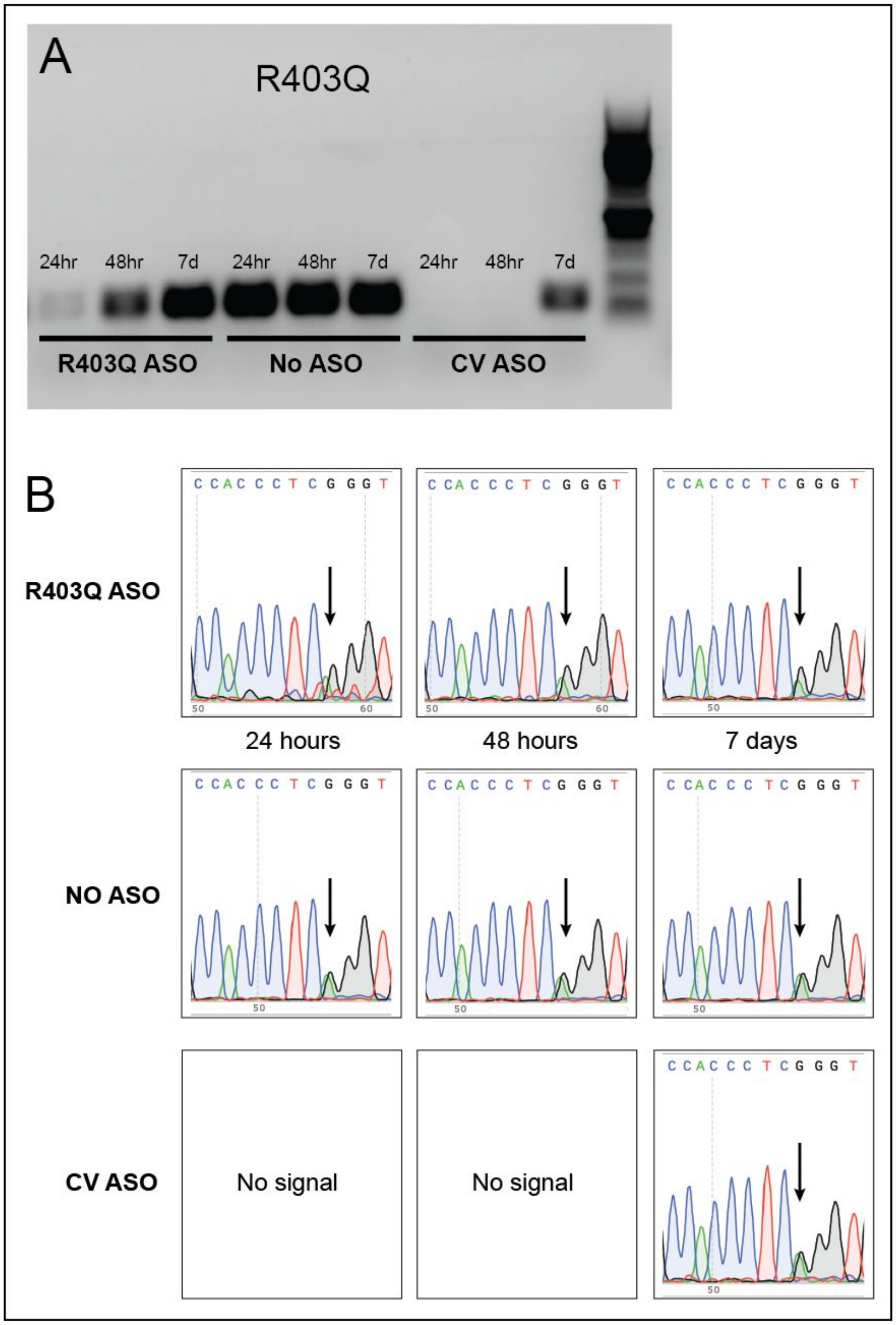


**S7 Fig: Sanger sequencing of R403Q locus in ASO-treated *MYH7*-R403Q hiPSC-CMs** Amplification of a 149bp region around the R403Q heterozygous variant in cDNA from 20-day old *MYH7*-R403Q hiPSC-CMs treated with either an R403Q-targeting ASO, no ASO, or an ASO designed to target a common variant in *MYH7* (CV ASO) which had previously been shown to knock down both the R403Q and R403R alleles. RNA was extracted 24 hours, 48 hours, or 7 days after transfection. In A) PCR of equal concentrations of cDNA from each sample revealed that *MYH7* amplification was high across all three timepoints in the No ASO control, while cells treated with the R403Q-targeting ASO showed diminished but non-zero amplification at 24 and 48 hours. However the cells treated with the CV ASO, previously shown to target both alleles, showed no amplification at 24 or 48 hours. In B) PCR fragments were sent out for Sanger sequencing. Traces from samples treated with the R403Q-targeting ASO show lower peaks from the R403Q allele (A) than the R403R allele (G). Peak heights are almost identical in the No ASO treated cells. The CV treated cells have no signal at hours 24 and 48 (consistent with the total or near-total knockdown seen in panel A) and no difference in peak height at 7 days, consistent with no discrimination between the two alleles. As the predicted Tm of the R403Q-targeting ASO is 38.2ºC, we do not expect that residual ASO extracted with our RNA should affect the results of these PCR reactions. PCR conditions: FWD Primer [AGCCAGACGGCACTGAAG], REV Primer [GGCATATATCACCTGCTGGA]. GoTaq Flexi DNA Polymerase protocol with cycling conditions 95C (2 min), 35-36 cycles of 95C (30 sec), 55.5C (30 sec), 72C (60 sec), followed by final 72C (5 min).

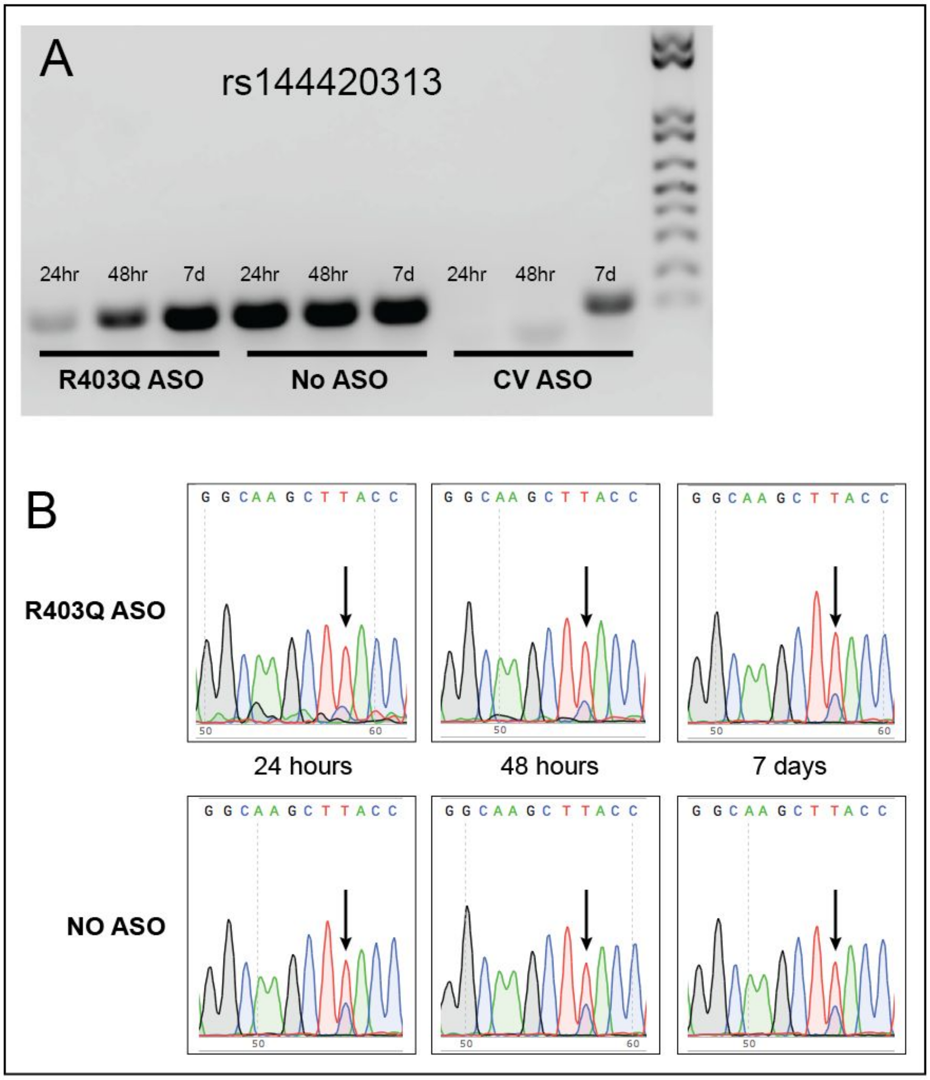


**S8 Fig: Sanger sequencing of rs144420313 locus in ASO-treated *MYH7*-R403Q hiPSC-CMs**

Amplification of a 149bp region around a heterozygous common variant (rs144420313) in cDNA from 20-day old *MYH7*-R403Q hiPSC-CMs treated with either an R403Q-targeting ASO, no ASO, or an ASO designed to target a common variant (rs144420313) in *MYH7* (CV ASO) which had previously been shown to knock down both the R403Q and R403R alleles. RNA was extracted 24 hours, 48 hours, or 7 days after transfection. In A) PCR of equal concentrations of cDNA from each sample revealed that *MYH7* amplification was high across all three timepoints in the No ASO control, while cells treated with the R403Q-targeting ASO showed diminished but non-zero amplification at 24 and 48 hours. However the cells treated with the CV ASO, previously shown to target both alleles, showed no amplification at 24 or 48 hours. In B) PCR fragments were sent out for Sanger sequencing. Peak heights in the No ASO treated cells are not identical for the two alleles, likely due to the high "T" peak right before the target SNP, where one of the possible bases is also a T. However, the relative height of the two traces do not change in this condition over time. Traces from samples treated with the R403Q-targeting ASO show lower peaks from the R403Q allele, here a C at the rs144420313 SNP, than the R403R allele (T) as compared to the No ASO control. As the R403Q-targeting ASO does not target this area of the transcript, we do not expect that residual R403Q-targeting ASO extracted with our RNA should affect the results of these PCR reactions. As the predicted Tm of the CV ASO is 36.5°C, we do not expect that residual ASO extracted with our RNA should affect the results of these PCR reactions either. PCR conditions: FWD Primer [AAGTTGCAAACCGAGAATGG], REV Primer [GCGTTCTTCGCCTTAACCTC]. GoTaq Flexi DNA Polymerase protocol with cycling conditions 95C (2 min), 33 cycles of 95C (30 sec), 55.5C (30 sec), 72C (30 sec), followed by final 72C (5 min).

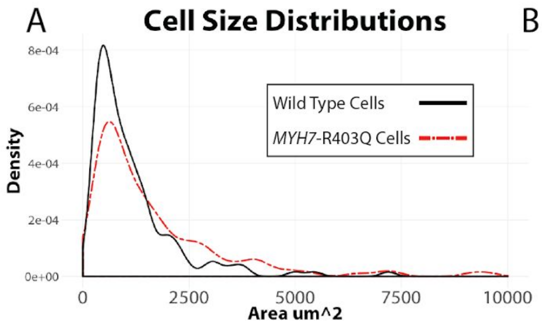


B

| Cell Line | Treatment | Mean (µm²) | Median (µm²) | n |
|---|---|---|---|---|
| **Wild Type** | NA | 1129 | 808 | 150 |
| **MYH7-R403Q** | NA | 1654 | 1076 | 147 |
| **MYH7-R403Q** | Transfection Reagent Only | 1620 | 1077 | 144 |
| **MYH7-R403Q** | R403Q-ASO + Transfection Reagent | 1753 | 916 | 146 |

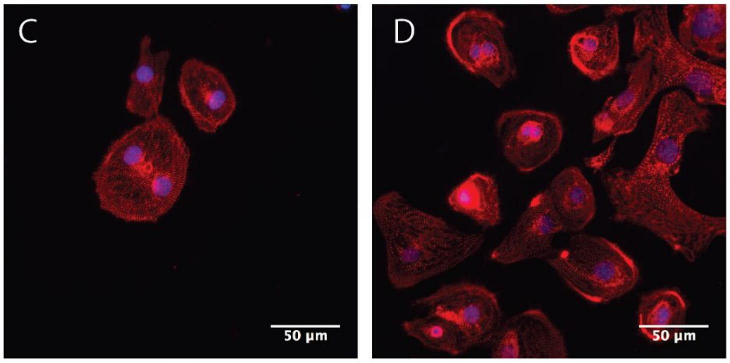


Wild Type Cells *MYH7*-R403Q Cells

**S9 Fig: Additional cell-size phenotyping measurements** A) Untreated *MYH7*-R403Q hiPSC-CMs fixed and stained at Day 30 showed an overall increase in cell population size as compared to a wild-type cell line (SCVI273). *MYH7*-R403Q hiPSC-CM median size = 1076µm$^2$, n = 147, vs Wild Type control cell line, median size = 808µm$^2$ , n = 150 , p=0.002207, Wilcoxon rank-sum test without continuity correction. All cells with sizes above 10,000µm$^2$ were removed prior to analysis. Plotting bandwidth adjustment = 0.8. B) Mean and median cell sizes for wild type or *MYH7*-R403Q cells treated with or without ASOs during differentiation. C&D) Representative immunofluorescent images. Blue is DAPI and red is α-actinin.

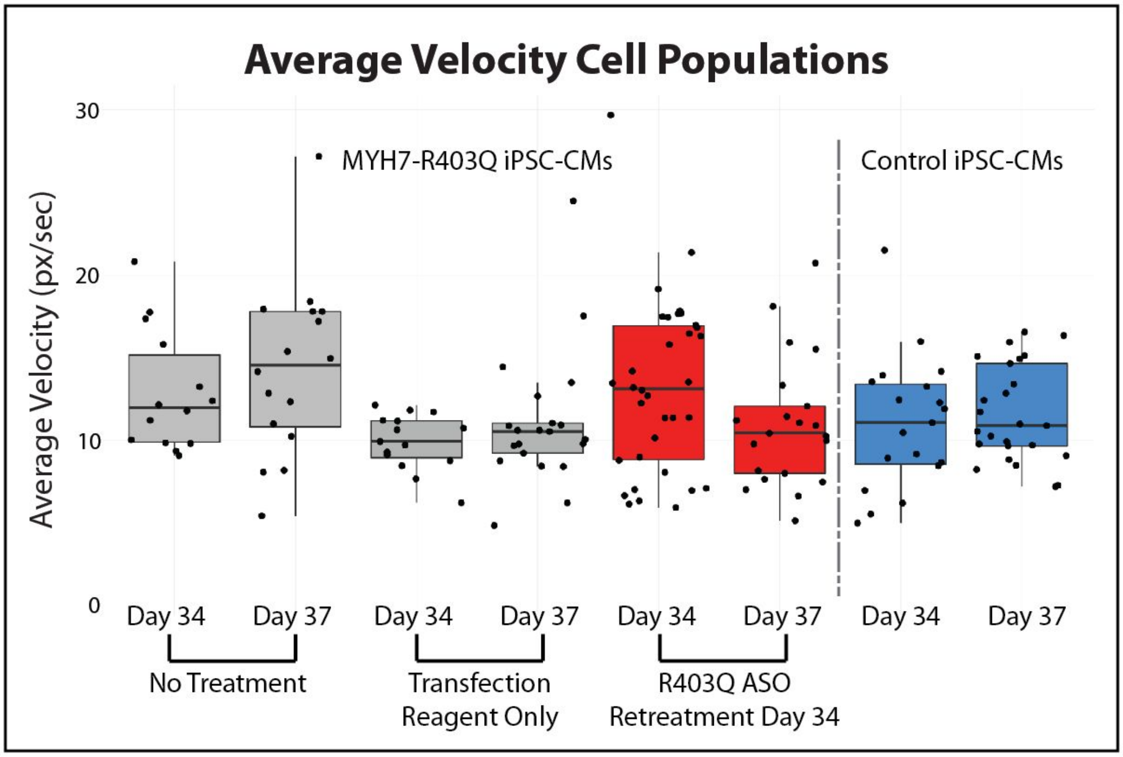


**S10 Fig: Additional cell velocity measurements** Plots of all imaged, beating, analyzed cells at days 34 and 37. Each point represents the average contractile velocity of a single cell over a 15 second video measured in pixels per second. Over the three day time period, *MYH7*-R403Q cells that were either untreated or treated with transfection reagent alone (gray) showed no change in their average contractile velocity. Wild-type controls cells additionally showed no change (blue). Cells retreated with the R403Q-targeting ASO showed a decrease in the population contractile velocity (red, Day 34 vs Day 37 p-value = 0.08083, two-sided t-test & p-value=0.04042, one-sided t-test.). Untreated *MYH7*-R403Q cells, Day 34 average: 12.9 px/sec, n = 14. Untreated *MYH7*-R403Q cells, Day 37 average: 14.3 px/sec, n = 16. Day 34 vs Day 37 p-value = 0.3988, two-sided t-test. Transfection-reagent-only treated *MYH7*-R403Q cells, Day 34 average: 9.9 px/sec, n = 15. Transfection-reagent-only treated *MYH7*-R403Q cells, Day 37 average: 11.1 px/sec, n = 21. Day 34 vs Day 37 p-value = 0.2507, two-sided t-test. R403Q-ASO treated *MYH7*-R403Q cells, Day 34 average: 13.2 px/sec, n = 34. R403Q-ASO retreated *MYH7*-R403Q cells,

Day 37 average: 11.0 px/sec, n = 21. Day 34 vs Day 37 p-value = 0.08083, two-sided t-test. Wild-Type Control cells, Day 34 average: 11.0 px/sec, n = 19. Wild-Type Control cells, Day 37 average: 11.6 px/sec, n = 25. Day 34 vs Day 37 p-value = 0.6081, two-sided t-test. Untreated *MYH7*-R403Q hiPSC-CMs show a slight increase in contractile velocity versus wild-type control cells at Day 37 (p-value = 0.0742, two-sided t-test & p-value = 0.0371, one-sided t-test).

| Target Gene | Primer Assay |
|---|---|
| EEF1A2 | Hs.PT.58.3514123 |
| MYH6 | Hs.PT.58.2106207 |
| MYH7 | Hs.PT.58.14589334 |
| NPPA | Hs.PT.58.4259173 |
| NPPB | HS.PT.58.19450190 |

**S2 Table: IDT qPCR Primers** qPCR Primers used to measure total gene expression. All primers were ordered from Integrated DNA Technologies. qPCR was performed using IDT's PrimeTime Master Mix using their recommended protocol on a ViiA 7 Real-Time PCR System (Thermo Fisher Scientific).

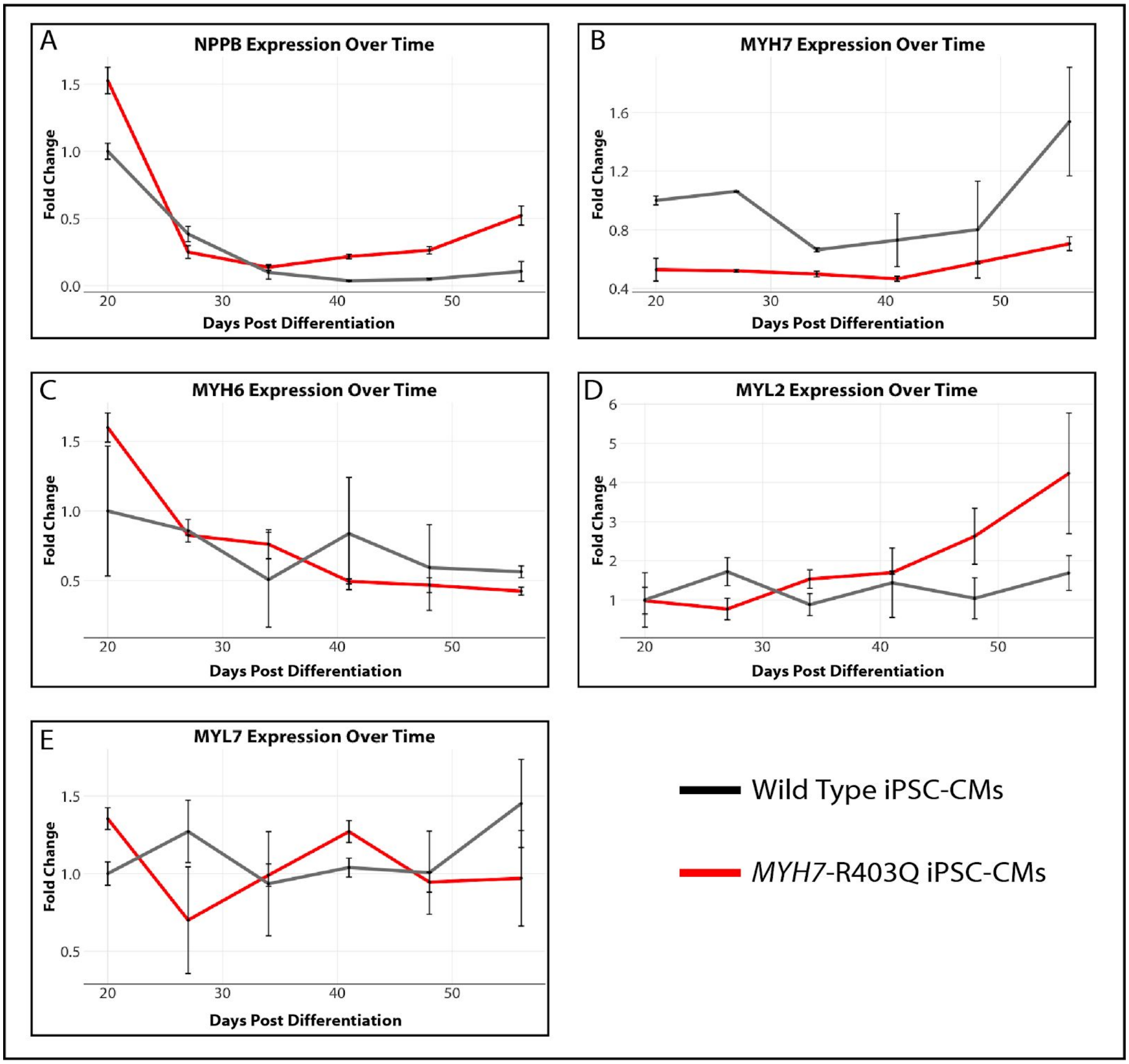


**S11 Fig: HCM-Associated RNA Expression Day 20 to Day 56 in differentiating hiPSC-CMs.** RNA expression levels of *NPPB*, *MYH7*, *MYH6*, *MYL2*, and *MYL7* in differentiating wild-type hiPSC-CMs (Control cell line SCVI34 obtained from the Stanford Cardiovascular Institute Biobank) or *MYH7*-R403Q HCM hiPSCs every seven days between Day 20 and Day 56 of differentiation. Each time point represents two biological replicate samples. Error bars are standard deviation of these two samples. All values normalized to SCVI34 wild type control cell line Day 20 values..
